# Impaired differential learning of fear versus safety signs in obsessive-compulsive disorder

**DOI:** 10.1101/2021.01.27.428235

**Authors:** Björn Elsner, Benedikt Reuter, Mahboba Said, Clas Linnman, Norbert Kathmann, Jan-Carl Beucke

## Abstract

Pavlovian learning mechanisms are of great importance both for models of psychiatric disorders and treatment approaches, but understudied in obsessive-compulsive disorder (OCD). Using an established Pavlovian fear conditioning and reversal procedure, we studied skin conductance responses (SCRs) in 41 patients with OCD and in 32 matched healthy control participants. Within both groups, fear acquisition and reversal effects were evident. When comparing groups, patients showed impaired differential learning of threatening and safe stimuli, consistent with previous research. In contrast to prior findings, differential learning impairments were restricted to fear acquisition, and not observed in the reversal stage of the experiment. As previous and present fear reversal experiments in OCD differed in the use of color coding to facilitate stimulus discrimination, the studies converge to suggest that differential learning of threatening versus safe stimuli is impaired in OCD, but manifests itself differently depending on the difficulty of the association to be learned. When supported by the addition of color, patients with OCD previously appeared to acquire an association early but failed to reverse it according to changed contingencies. In absence of such color coding of stimuli, our data suggest that patients with OCD already show differential learning impairments during fear acquisition, which may relate to findings of altered coping with uncertainty previously observed in OCD. Impaired differential learning of threatening versus safe stimuli should be studied further in OCD, in order to determine whether impairments in differential learning predict CBT treatment outcomes in patients, and whether they are etiologically relevant for OCD.

## Introduction

Pavlovian fear conditioning mechanisms are commonly used to explain a variety of phenomena in psychopathology. They are particularly highlighted by models intending to explain the etiology and maintenance of fear symptoms (Mineka & Oehlberg, 2008; Mowrer, 1939), and are viewed to be fundamental to the action of cognitive behavioral therapy (CBT). Therapists may educate patients that they have learned to fear certain objects or situations (abnormal fear acquisition), and that the goal of the therapy is to modify this acquired fear response (by fear extinction or reversal). Indeed, core elements of CBT such as exposure with response prevention (ERP) to some extent resemble components of experimental laboratory trainings of fear reversal or extinction, and their application is based on consistent evidence of efficacy. In obsessive-compulsive disorder (OCD), for example, ERP effectiveness has been demonstrated using randomized controlled trials (Olatunji, Davis, Powers, & Smits, 2013; Öst, Havnen, Hansen, & Kvale, 2015) as well as under routine care conditions (Elsner et al., 2020; Hansen, Kvale, Hagen, Havnen, & Öst, 2018). Nevertheless, further improvements in treatment efficacy are necessary and will require the introduction of new therapeutic approaches, as well as optimization of the currently existing procedures (Mataix-Cols, Fernandez De La Cruz, & Rück, 2019). In clinical practice, ERP sessions are designed, adjusted and tailored for the single case at hand, particularly in the case of OCD and its known symptom heterogeneity (Mataix-Cols, Rosario-Campos, & Leckman, 2005). It is thus of crucial importance to understand deficient fear processing mechanisms and whether they concern fear acquisition, reversal or retention of extinction processes. The precise identification of the stages or phases where fear learning is altered may improve treatment. This may be of particular importance for treatment of OCD, as a substantial portion of patients with OCD does not benefit from CBT (Mataix-Cols et al., 2019; Öst et al., 2015; Taylor, Abramowitz, & Mckay, 2012).

Pavlovian learning studies in anxiety disorders typically show heightened fear responses to safe stimuli during fear acquisition, and exaggerated responses to the previously threatening, but now safe stimuli during extinction in patients (Duits et al., 2015). Only a very small number of studies investigated Pavlovian fear learning deficits in OCD to date (Apergis-Schoute et al., 2017; Mclaughlin et al., 2015; Milad et al., 2013; Nanbu et al., 2010). The existing studies point to unaltered fear acquisition and extinction (Mclaughlin et al., 2015; Milad et al., 2013; Nanbu et al., 2010), but impaired extinction recall (Mclaughlin et al., 2015; Milad et al., 2013), and reduced differentiation of safe and threatening stimuli during late phases of fear acquisition and reversal (Apergis-Schoute et al., 2017) in OCD. Whereas traditional Pavlovian learning paradigms implement pauses of several hours or even days between fear acquisition and extinction recall (Mclaughlin et al., 2015; Milad et al., 2013), the fear reversal paradigm considers that fear responses need to be flexibly and rapidly adjusted to changing contingencies. It involves a transient acquisition and reversal of fear within the same, single experimental session. In the acquisition stage, two facial cues are presented, and paired with an aversive stimulus (CS+) or no outcome (CS-). Later, in the reversal stage of the experiment, the previously aversive cue becomes the safe stimulus (new CS-), whereas the previously safe stimulus becomes the threatening stimulus now associated with the aversive outcome (new CS+; Schiller, Levy, Niv, Ledoux, & Phelps, 2008). Acquisition and reversal stages are further subdivided in early and late phases. In a study of healthy participants, Schiller et al. (2008) demonstrated significantly greater skin conductance responses (SCRs) to the CS+ compared to the CS-during acquisition, and significant SCR differentiation between the new CS+ and the new CS-during reversal, suggesting that fear learning can be successfully acquired and reversed. Notably, the awareness of contingency reversal can differ between clinical and control groups (Apergis-Schoute et al., 2017) and thus needs to be carefully considered in group comparisons of fear reversal responses. The fear reversal paradigm may be of particular relevance for patients with OCD, as impairments in the context of operant conditioning are consistently found when stimulus contingencies are reversed (Chamberlain et al., 2008; Remijnse et al., 2006; Valerius, Lumpp, Kuelz, Freyer, & Voderholzer, 2008). Using a modified version of the fear reversal experiment in which the two facial cues were overlaid with two different colors, a previous study in OCD and healthy control participants (Apergis-Schoute et al., 2017) showed significantly impaired differentiation of CS+ and CS-throughout the entire experiment for patients with OCD, which were found in late but not in early acquisition and reversal phases, respectively. Within both groups, significant differentiation of CS+ and CS-was evident during fear acquisition, however during reversal, no such effects were observed in patients, in contrast to significant differential learning in healthy controls (Apergis-Schoute et al., 2017).

Action of antidepressant medication represents a relevant confounder in the study of fear responses in patient populations. Studies in rats suggest that chronic citalopram administration impairs the acquisition of fear conditioning (Burghardt, Sullivan, Mcewen, Gorman, & Ledoux, 2004), and chronic antidepressant treatment impairs the acquisition of extinction (Burghardt, Sigurdsson, Gorman, Mcewen, & Ledoux, 2013), respectively. Further, studies in humans revealed that serotonergic depletion specifically impairs classical fear conditioning (Hensman, Guimarães, Wang, & Deakin, 1991; Hindi Attar, Finckh, & Büchel, 2012) and fear reversal learning (Kanen et al., 2020). Moreover, escitalopram seems to facilitate extinction learning (Bui et al., 2013), and finally, antidepressant medication emerged as a key variable when comparing brain metrics between patients with OCD and healthy subjects in a recent mega-analysis in OCD (Boedhoe et al., 2017).

The present study sought to investigate fear reversal learning in patients with OCD in absence of the artifacts and possible claustrophobic tendencies that may be induced by simultaneous functional magnetic resonance imaging recordings present in previous studies (Apergis-Schoute et al., 2017; Milad et al., 2013). We further carefully considered antidepressant medication effects, and controlled for putative differences in contingency reversal awareness when comparing fear reversal responses between patients and controls. The addition of color to facial stimuli, which likely facilitates stimulus discrimination, was omitted. Three questions were addressed: First, we hypothesized that previously reported fear reversal effects (Schiller et al., 2008) are replicable. Second, we tested the hypothesis that patients with OCD display reduced differential fear learning of threatening versus safe stimuli. We expected such reductions to be most pronounced during late acquisition and reversal phases, as indicated previously (Apergis-Schoute et al., 2017). Finally, we assessed the effect of antidepressant medication by comparing fear acquisition and reversal responses between medicated and unmedicated patients.

## Method

### Experimental Procedure and Data Acquisition

We implemented the experimental setting of the previous fear reversal study by Schiller et al. (2008). A mild electrical stimulation served as unconditioned stimulus (US). The amperage of shock level was individually selected prior to the experiment so that stimulation was “unpleasant but not painful”. The shock (bursts of 50 ms) was delivered to the dominant wrist. As previously (Schiller et al., 2008), two mildly angry faces from the Ekman series (Ekman & Friesen, 1976) served as conditioned stimuli (CS) and were presented for four seconds per trial with an intertrial interval (ITI) of twelve seconds.

The experiment involves two *stages*: During *acquisition* (30 trials), face A (CS+) was presented in 18 trials and paired with the shock in one-third of these trials (six trials; Schiller et al., 2008). Face B (CS-) was presented in twelve trials and was never paired with the shock. During *reversal* (40 trials), contingencies were reversed without announcement: face B (new CS+) was presented in 24 trials and now paired with the shock in one-third of these trials (eight trials) while face A (new CS-) was presented in 16 trials and now no longer shocked (compare Schiller et al., 2008, Figure 1). To avoid order effects, the array of trials varied regarding the occurrence of first shock delivery across three different versions. In all versions, there was no reinforcement of the first CS+, and there were no consecutive reinforced trials. Further, the assignment of faces into CS+ and CS-was counterbalanced across all subjects. We divided the acquisition and reversal stages in early and late trials (*phase*) in order to observe contingency learning over time (Schiller et al., 2008).

**Figure 1.**
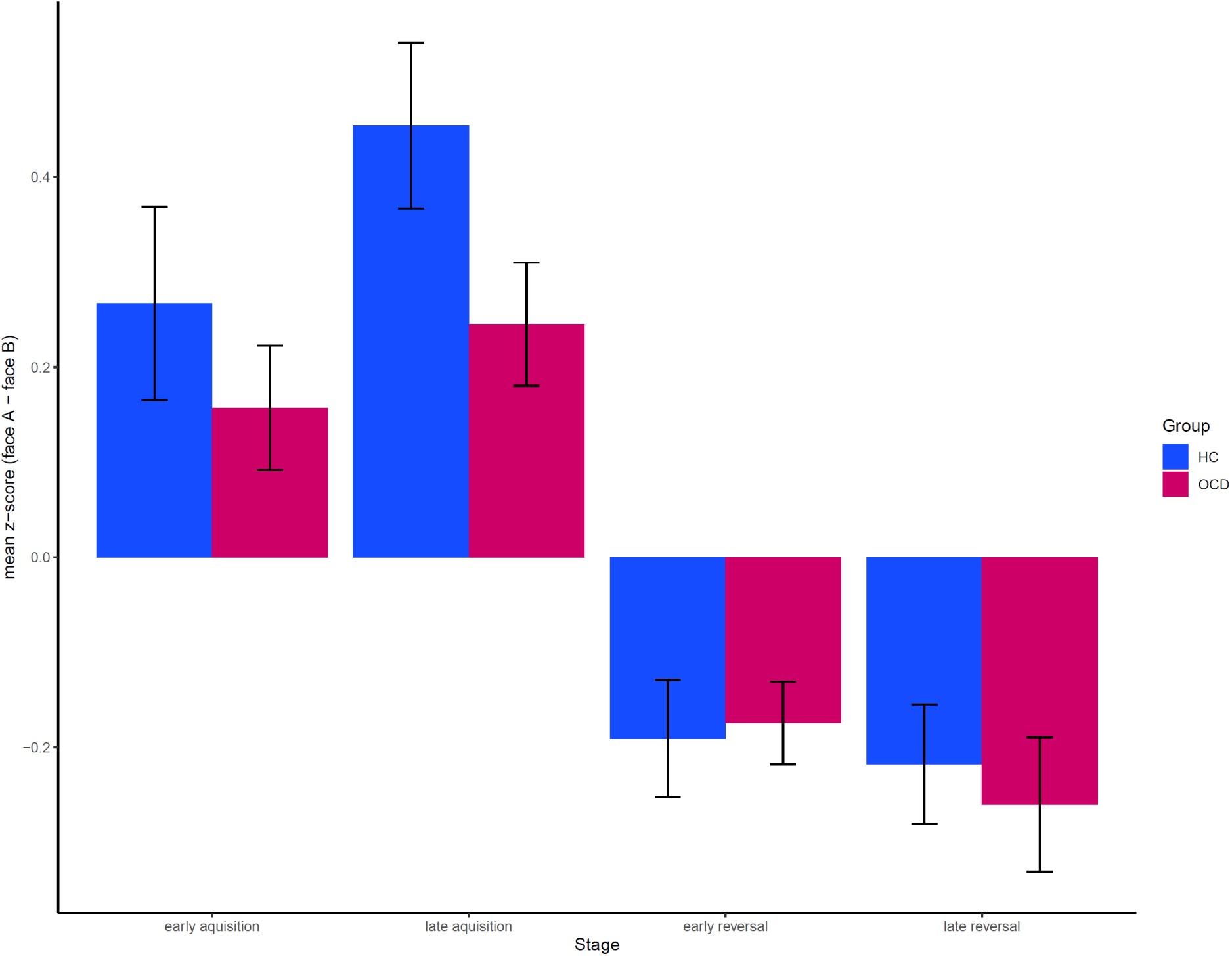
Differential responding in z-transformed SCR to face A (CS+ in acquisition, CS-in reversal) vs. face B (CS+ in reversal, CS-in acquisition), indicating successful fear acquisition and reversal in both groups as described in the original study (compare to Schiller et al, 2008, Fig. 2A), as well as group differences in fear acquisition. Error bars indicate ±1 standard errors of mean differences.

Skin conductance response (SCR) levels during anticipation of a potential shock (0.5 – 4.5 s after trial onset) served as the dependent variable. SCR were measured using Ag-AgCl electrodes (1 cm^2^) which were attached hypothenar to the non-dominant hand.

Participants were instructed to pay attention to the relationship between the faces and the occurrence of the shocks. After the experiment we assessed explicit contingency awareness by asking what participants noticed regarding the relationship between faces and shocks. Thus, a binary variable indicating awareness of contingency reversal was derived and used for further analysis.

The study was approved by the local review board of Humboldt-Universität zu Berlin (protocol numbers 2016-29 and 2019-05) and met the standards of the revised Declaration of Helsinki. All participants provided written informed consent and received 10 EUR per hour of participation.

### Psychological Assessment

Immediately before the experiment, a clinical psychologist conducted the German version of the Yale-Brown Obsessive-Compulsive Scale interview (Y-BOCS; Goodman et al., 1989a; Goodman et al., 1989b; Hand & Büttner-Westphal, 1991) and the Montgomery Åsberg Depression Rating Scale (MADRS; Montgomery & Åsberg, 1979) with all OCD patients. After termination of the experimental procedure, all participants completed several psychological tests and questionnaires. They included the Wortschatztest (treasury of words test; Schmidt & Metzler, 1992) in order to estimate verbal intelligence, and a socio-economic status scale (Lampert & Kroll, 2009). Moreover, the Beck Depression Inventory II (BDI-II; Beck, Steer, & Brown, 1996), the Obsessive Compulsive Inventory-Revised (OCI-R; Foa et al., 2002) and the State-Trait Anxiety Inventory (STAI-X; Spielberger, Gorsuch, & Lushene, 1970) were used in order to assess symptom severity of depressive, OCD, and anxiety symptoms, respectively.

### Participants

Eighty-two subjects participated in the experiment. Data from two participants were discarded due to technical problems during data recording. Further, visual data inspection revealed that seven participants (three OCD) did not show any SCR to shocked trials (US). In line with recent recommendations (Lonsdorf et al., 2019), they were classified as non-responders and not considered for further analysis.

The final sample comprised 41 patients (25 female; mean age = 29.93 years; SD = 8.54) with a primary diagnosis of OCD and 32 matched healthy controls (HC, 17 female; mean age = 28.97 years; SD = 7.49). All patients were registered on a waiting list for cognitive behavioral therapy at the outpatient unit for psychotherapy at Humboldt-Universität zu Berlin.

Patients and control participants did not differ significantly in demographic or basic experimental variables, however, OCD patients scored significantly higher on the clinical assessments, see Table 1. In the group of OCD patients, comorbid diagnoses of mental disorders were anxiety disorders (n = 6), somatoform disorder (n = 1), and depressive disorder (n = 20). Twenty-eight patients were free of psychotropic medication as defined by an a priori washout criterion requiring no intake of any psychotropic medication for at least three months prior to the experiment. Thirteen patients did not fulfill this criterion due to treatment with antidepressants (n = 3 escitalopram, n = 2 citalopram, n = 4 paroxetine, n = 3 sertraline, n = 1 hypericum).

**Table 1.** Group differences in demographic, basic experimental and clinical variables

| *Variable* | *HC* | *OCD* | *Test* |
| --- | --- | --- | --- |
| Gender | 17 f / 15 m<br>*M(SD)* | 25 f / 16 m<br>*M(SD)* | $\chi^2(1) = .45, p = .501$ |
| Age (yrs) | 29.0 (7.5) | 29.9 (8.5) | $t(71) = -0.50, p = .617$ |
| Verbal intelligence (IQ) | 106.8 (8.4) | 104.2 (9.6) | $t(71) = 1.21, p = .230$ |
| Socio-economic status | 11.1 (3.0) | 9.8 (2.9) | $t(71) = 1.89, p = .063$ |
| Impedance stimulation electrode (kΩ) | 40.2 (29.3) | 46.4 (27.7) | $t(71) = -0.92, p = .361$ |
| Selected amperage of shock level (mA) | 1.6 (1.8) | 1.8 (1.3) | $t(71) = -0.33, p = .746$ |
| SCR to electrical stimulation (z) | 1.1 (0.5) | 1.3 (0.6) | $t(71) = -0.86, p = .392$ |
| OCI-R | 5.1 (5.0) | 27.0 (12.8) | $t(54.53) = -9.14, p < .001$ |
| BDI-II | 3.6 (5.7) | 13.3 (10.2) | $t(64.85) = -4.81, p < .001$ |
| STAI-X1 | 31.4 (5.1) | 41.2 (10.7) | $t(60.29) = -4.82, p < .001$ |
| STAI-X2 | 32.4 (6.2) | 50.3 (12.5) | $t(61.41) = -9.14, p < .001$ |
| Y-BOCS | - | 22.3 (4.5) | - |
| MADRS | - | 7.54 (8.0) | - |

*Note.* OCI-R = Obsessive Compulsive Inventory - Revised; BDI-II = Beck Depression Inventory II; STAI-
X = State-Trait Anxiety Inventory; Y-BOCS = Yale-Brown Obsessive-Compulsive Scale interview score;
MADRS = Montgomery Åsberg Depression Rating Scale

### Data Analysis

Psychophysiological raw data were recorded using Brain Vision Recorder, exported with Brain Vision Analyzer (Version 2.1), and finally transferred to be analyzed in Matlab (Version R2016a). Using the Ledalab toolbox (Version 3.4.9) data were downsampled from 1000 Hz to 100 Hz. A Butterworth lowpass filter was applied to the data. In order to dissociate continuous phasic and tonic activity a continuous decomposition analysis (Benedek & Kaernbach, 2010) was used. SCRs were z-transformed in order to normalize data, also accounting for responses to the shock. Thus, mean scores may have negative values but were consistently positive prior to z-transformation (see Figure 2). A SCR-window of 0.5 to 4.5 s was exported. Data epochs below the minimum amplitude threshold of 0.02 μS were coded as zero. The key outcome measure was the mean of above-threshold SCR-amplitudes within the response window calculated across all trials of a condition. Given the absence of MRI measurements, trials reinforced by electrical stimulation were also included in our analysis, in contrast to the previous studies (Apergis-Schoute et al., 2017; Schiller et al., 2008). Moreover, we removed trials before the first shock in both stages (the orienting response; Cohen, 1993), and the first CS-trial after the shock during reversal, as participants had no knowledge of the current contingencies at that particular part of the experiment. Individual data were exported to SPSS (Version 23) for statistical analysis.

**Figure 2.**
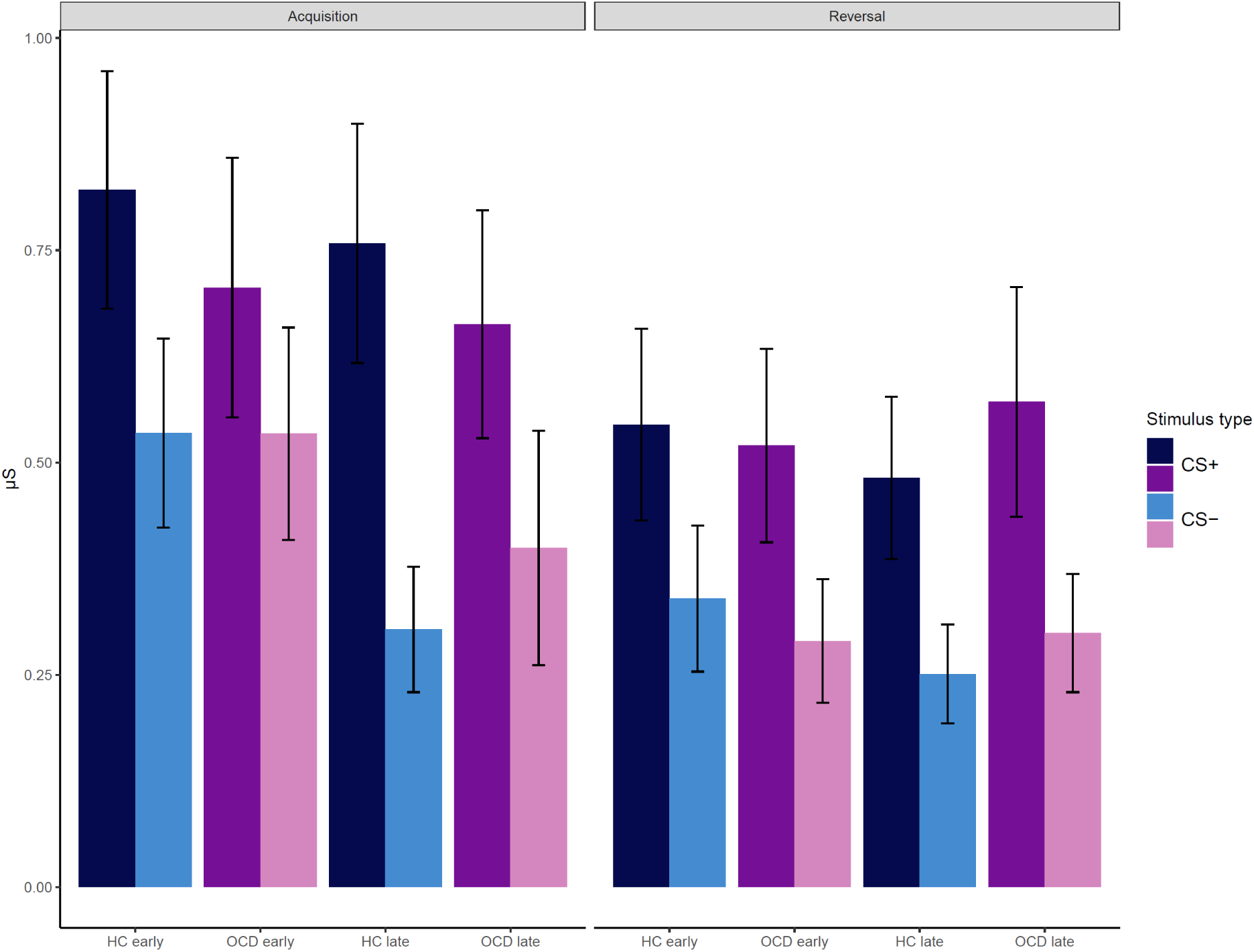
Mean SCR values for each condition, plotted separately for group (HC and OCD) and stimulus type (CS+ and CS-). Error bars indicate standard errors.

### Statistical Analysis

In order to test demographic and questionnaire variables for group differences, independent t-tests were conducted to compare means, and a Pearson Chi-Square test was used to compare nominal data.

We analyzed mean SCR-amplitudes using a 2×2×2×2×2 repeated measures-ANOVA with *stage* (acquisition vs. reversal), *phase* (early vs. late) and *stimulus* (CS+ vs. CS-) as within subject factors, and *group* (HC vs. OCD) and contingency reversal *awareness* (aware vs. unaware) as between subjects factors. Post-hoc t-tests were conducted to explain significant interactions. Further, in order to test for effects of antidepressant medication, we compared SC responses between unmedicated (n = 28) and medicated (n = 13) patients directly by repeating the ANOVA after replacing the *group* (HC vs. OCD) factor with a *medication* (unmedicated patients vs. medicated patients) factor.

In an exploratory analysis, we correlated scores on the Y-BOCS scale with differential skin conductance responses from the acquisition and the reversal stage within the OCD group. The significance level was set at α = .05 for all tests.

## Results

Effects of fear learning and fear reversal were observed in the combined group of subjects. All main and interaction effects of the ANOVA on SCR amplitudes are displayed in Table 2. Figure 1 displays differential responding as z-transformed SCRs to face A versus face B for both groups. Figure 2 indicates mean scores of SCR before z-transformation for each condition, plotted separately for group and stimulus type.

**Table 2.** ANOVA with repeated measures: Main and interaction effects including post-hoc t-tests

| | F (1,69) | p | partial $\eta^2$ |
| --- | --- | --- | --- |
| ***Replication of fear reversal effects*** |  |  |  |
| Stage (acquisition vs. reversal) | = 22.45 | < .001 | .245 |
| Phase (early vs. late) | = 3.77 | = .056 | .052 |
| Stimulus (CS+ vs. CS-) | = 39.13 | < .001 | .362 |
| Stage x phase | = 0.67 | = .414 | .010 |
| Stage x stimulus | = 1.57 | = .215 | .022 |
| Phase x stimulus | = 3.85 | = .054 | .053 |
| Stage x phase x stimulus | = 0.30 | = .585 | .004 |
| ***Group differences*** |  |  |  |
| Group (HC vs. OCD) | = 0.25 | = .621 | .004 |
| Group x stage | = 4.54 | = .037 | .062 |
| - for acquisition: HC > OCD | t(71) = 2.16 | = .034 |  |
| Group x phase | = 3.80 | = .055 | .052 |
| - for early: HC > OCD | t(71) = 1.99 | = .050 |  |
| Group x stimulus | = 0.57 | = .453 | .008 |
| Group x stage x phase | = 0.05 | = .833 | .001 |
| Group x stage x stimulus | = 5.48 | = .022 | .074 |
| - for CS+ in acquisition: HC > OCD | t(71) = 2.64 | = .010 |  |
| Group x phase x stimulus | = 0.01 | = .926 | .000 |
| Group x stage x phase x stimulus | = 0.18 | = .675 | .003 |
| ***Reversal awareness differences*** |  |  |  |
| Awareness (aware vs. unaware) | = 0.15 | = .703 | .002 |
| Awareness x group | = 0.64 | = .425 | .009 |
| Awareness x stage | = 0.20 | = .654 | .003 |
| Awareness x stage x group | = 0.05 | = .830 | .001 |
| Awareness x phase | = 0.16 | = .693 | .002 |
| Awareness x phase x group | = 0.67 | = .418 | .010 |
| Awareness x stimulus | = 4.19 | = .044 | .057 |
| - for CS-: unaware > aware | t(71) = 2.66 | = .010 |  |
| Awareness x stimulus x group | = 0.05 | = .833 | .001 |
| Awareness x stage x phase | = 0.16 | = .689 | .002 |
| Awareness x stage x phase x group | =0.19 | = .661 | .003 |
| Awareness x stage x stimulus | = 0.72 | = .400 | .010 |
| Awareness x stage x stimulus x group | = 2.36 | = .129 | .033 |
| Awareness x phase x stimulus | = 0.15 | = .698 | .002 |
| Awareness x phase x stimulus x group | = 0.06 | = .816 | .001 |
| Awareness x stage x phase x stimulus | = 0.77 | = .384 | .011 |
| Awareness x stage x phase x stimulus x<br>group | = 1.40 | = .241 | .020 |

### Fear reversal effects

The ANOVA revealed a significant main effect of *stage*, indicating stronger SCR for acquisition (M = −0.197, SD = 0.17) than for reversal (M = −0.331, SD = 0.15; Table 2). Further, the main effect of *stimulus* (CS+ vs. CS-) was significant, indicating stronger SCR for CS+ (M = −0.143, SD = 0.21) than for CS-(M = −0.384, SD = 0.13).

### Group differences

The interaction between *group* and *stage* was significant (Table 2 and Figure 1). Post-hoc tests revealed significantly stronger SCRs during acquisition for healthy controls (M = −0.149, SD = 0.18) than for OCD patients (M = −0.234, SD = 0.16), t(71) = 2.16, p = .034. No significant group difference could be observed during reversal, t (71) = −1.09, p = .278. We also observed a significant three-way interaction between *group, stage* and *stimulus* (Table 2). Post-hoc t-tests revealed significantly stronger SCRs for healthy control participants (M = 0.031, SD = 0.30) than for OCD patients (M = −0.134, SD = 0.23) specifically for CS+ during acquisition, t(71) = 2.64, p = .010 (Figure 2). Finally, the interaction between *group* and *phase* also reached trend level (Table 2). Post-hoc t-tests indicated a tendency towards stronger responses for healthy controls in early trials, (M = −0.201, SD = 0.17) than for OCD patients (M = −0.268, SD = 0.12), t(71) = 1.99, p = .050. The interaction between *group* and *stimulus* did not yield significance (p = .453), indicating no general difference between healthy control participants and OCD patients regarding their differential SCRs.

A significant interaction between contingency reversal *awareness* and *stimulus* (Table 2) was observed. Post-hoc tests indicated that participants who were not aware of the reversal showed stronger responses to the CS-(M = −0.325, SD = 0.11) compared to participants who reported contingency reversal awareness (M = −0.410, SD = 0.13), t(71) = 2.66, p = .010. The interaction between *awareness* and *group* was not significant (Table 2). Direct comparison of contingency reversal awareness between groups (that is, irrespective of SCR) revealed no significant differences in awareness (OCD = 27, HC = 24) and unawareness (OCD = 14, HC = 8) of the contingency reversal, χ^2^(1) = 0.71, p = .398.

### Medication effects

ANOVA was repeated to compare effects in unmedicated (n = 28) versus medicated (n =13) OCD patients (Supplementary Table 3). Unmedicated patients did not differ from medicated patients regarding demographic and basic experimental variables (Supplementary Table 2). The main effects for *stage* and for *stimulus* remained significant. However, no significant interaction with *medication* (unmedicated vs. medicated) was observed, indicating no influence of medication on *stage, phase, stimulus*, contingency reversal *awareness*, or their interactions, respectively.

### Exploratory analyses

For the group of patients with OCD, no significant correlations were observed between Y-BOCS scores and the difference score between CS+ and CS-(acquisition: r = .287, p = .079; reversal: r = .048, p = .766).

## Discussion

The present study investigated skin conductance responses (SCRs) of patients with OCD as compared to healthy individuals in a fear reversal paradigm. In accordance with previous findings (Schiller et al., 2008), SCRs indicated a distinct pattern of fear acquisition and reversal effects across both groups. Compared with healthy control participants, OCD patients showed impaired differential learning of threatening versus safe stimuli during fear acquisition, explained by a significantly smaller SCR to CS+ in this stage of the experiment. While these results converge with previous findings of altered differential learning of threatening versus safe stimuli in OCD, impaired differential learning was not observed in the fear reversal stage, in contrast to previous research (Apergis-Schoute et al., 2017). These discrepancies between prior and present fear reversal studies in OCD may be explained by differences in the degree of difficulty between studies regarding the associations to be learned. When stimulus discrimination is supported by color coding, patients with OCD may acquire the association but fail to reverse it, whereas the absence of such facilitation reveals that patients are already impaired in differential learning of threat versus safety when forming associations in the initial process of fear acquisition. However, this study was not designed to directly compare the effects of color on discrimination learning.

Irrespective of group differences, the present results are consistent with the existing literature in demonstrating effects of fear acquisition and reversal. Although the replicability of this pattern initially described by Schiller et al. (2008) had previously been indicated by Apergis-Schoute et al. (2017), modification of the experimental procedure (addition of colors to the two facial stimuli serving as CS+ and CS-) has substantially facilitated stimulus discrimination in that study. In the present study, we demonstrate that fear reversal effects as reported by Schiller et al. (2008) can be replicated independent of such facilitation. The relevance of this aspect is highlighted by the significant post-session contingency reversal awareness by stimulus interaction observed, because facilitation of stimulus discrimination likely leads to higher contingency reversal awareness, which in turn affects differential learning, as the interaction was explained by the fact that participants who were unaware of the reversal displayed higher SC responses to safe (CS-) stimuli. In this manner, facilitating the discrimination of conditioned stimuli may significantly impact differential learning of CS+ versus CS-and reversal learning effects. The present replication of the fear reversal effect without additional color coding (Schiller et al., 2008) is therefore non-trivial and corroborates the robustness of fear reversal effects and the suitability of the paradigm for future research.

Group differences in the present study emerged as significant *group by stage* and *group by stage by stimulus* interactions, respectively, indicating impaired differential learning of threatening versus safe stimuli during fear acquisition in patients with OCD, which was explained by reduced SCR to the CS+. This confirms earlier evidence of impaired differential learning of threatening versus safe stimuli (Apergis-Schoute et al., 2017), but also deviates from previous results, as the differential learning deficit was restricted to the fear acquisition stage in the present, but was also found in late phases of both the acquisition and reversal stages in the previous study. Our finding also differs from results from other classical conditioning studies in OCD which neither found acquisition deficits for patients (Mclaughlin et al., 2015; Milad et al., 2013). Further, we did not observe globally altered learning in the form of a main effect of group in differential learning or a group by stimulus interaction, as reported previously (Apergis-Schoute et al., 2017). As confounding effects of antidepressant medication did not appear to impact the results, the discrepancy between the present and the previous results is likely best explained by differences in study design, experimental setup and analysis strategy (Supplementary Table 1). Among those differential aspects, the aforementioned use of colors facilitating stimulus discrimination holds potential to partially explain the deviating results, as the use of these additional cues impacts the degree of difficulty of the associations to be learned, and generates different levels of uncertainty or volatility in the context of associative learning between the two studies. More precisely, the addition of colors may cover or partially compensate a basic deficit in forming fear associations under uncertain conditions in patients with OCD, and may allow for acquisition of fear associations which patients later fail to reverse to changing contingencies. A more uncertain state in absence of color coding support appears to impair the process of fear acquisition already, which may in turn prevent observation of a deficit in fear reversal. A further notable difference among the two studies that may relate to deviating fear reversal results is that patients revealed significantly less contingency reversal awareness during the experiment compared to control participants previously (patients were found to have significantly less knowledge than controls; Apergis-Schoute et al., 2017), whereas no such group difference was observed in the present study.

Exploratory analyses testing for associations between fear acquisition impairments and OCD symptom severity did not reveal significant correlations, confirming the previous study (Apergis-Schoute et al., 2017). This suggests that the deficits are unrelated to current symptoms but may rather represent a state-independent impairment. Future longitudinal studies are needed to verify whether the observed fear learning deficit is predictive of ERP treatment responses and the clinical course of OCD. There is early evidence that therapy outcome can be predicted by SCRs in childhood OCD (Geller et al., 2019). It is tempting to speculate that the apparent symptom-independence could indicate that fear learning abnormalities represent a candidate endophenotype and may be associated with familial vulnerability for OCD. However, evidence regarding this aspect is lacking so far. Investigation of putative fear learning abnormalities in OCD-unaffected first-degree relatives or ideally monozygotic twin pairs discordant for OCD would be helpful to clarify the role of impaired differential Pavlovian learning of threat versus safety in the etiology of OCD.

As prior and present studies converge to suggest a basic deficit in differentiating threatening and safe stimuli in OCD, it is worth considering associations with other behavioral abnormalities in OCD. It seems plausible to assume that patients overgeneralize fear responses when stimulus contingencies are unclear. Such a tendency is known in panic disorder (Lissek et al., 2010) and generalized anxiety disorder (Lissek et al., 2014), and has also been associated with high levels of obsessive-compulsive traits and high threat estimation in the context of conditioned fear (Kaczkurkin & Lissek, 2013). Further, the differential learning deficit may relate to altered coping with uncertainty as indicated by a previous study where OCD patients showed a lack of differentiation between unpredictable and predictable threat stimuli in a cued picture paradigm in absence of symptom-relevant stimuli (Dieterich, Endrass, & Kathmann, 2017), possibly reflecting increased attentional responses to uncertainty. Taken together, patients with OCD appear to display altered processing and coping in volatile environments with uncertain contingencies, where they show impairments in differentiation of safe versus threatening cues.

In summary, the present findings reveal a basic Pavlovian learning deficit in OCD, as patients were characterized by impairments in differentiating threatening and safe stimuli during fear acquisition. Present results thereby confirm the core finding of impaired differential learning from a previous fear reversal study in patients with OCD (Apergis-Schoute et al., 2017). The allocation of this deficit in distinct stages or phases of the fear acquisition and reversal procedure may depend on experimental variables influencing contingency uncertainty and the difficulty of the association to be learned, as well as group differences in awareness of contingency reversal. Across patient and nonclinical groups, fear reversal revealed to be stable and reproducible, suggesting that the fear reversal procedure may be suitable for longitudinal study efforts. Future studies should address the value of the observed deficits in predicting CBT treatment response, and attempt to learn more about the role of Pavlovian fear acquisition impairments in the etiology of OCD.

## Acknowledgments

The authors assert that all procedures contributing to this work comply with the ethical standards of the relevant national and international committees on human experimentation and with the Helsinki Declaration of 1975, as revised in 2008. This research received no specific grant from any funding agency, commercial or not-for-profit sectors. The authors thank Sophie Harms for conducting pilot experiments, Rainer Kniesche for technical support, and Ulrike Lüken, Lauriane Tasiaux, Stefanie Kunas, Helena Köthner for collaboration and recruitment.

## Supplementary Tables

**Supplementary Table 1.** Overview of differences and similarities in critical experimental aspects and statistical analysis when comparing the original fear reversal study in healthy individuals (Schiller et al., 2008) to a previous study in OCD (Apergis-Schoute et al., 2017) and the present study, respectively.

|  | Schiller et al., 2008 | Apergis-Schoute et al., 2017 | Present Study |
| --- | --- | --- | --- |
| Availability of simultaneously recorded fMRI data | ✓ | ✓ | ✗ |
| Acquisition of SCR data in shielded room in absence of MR artifact | ✗ | ✗ | ✓ |
| Use of color on facial stimuli to facilitate discrimination learning | ✗ | ✓ | ✗ |
| Counterbalancing: Use of alternating versions of stimulus and shock delivery to avoid order effects | ✗ | ✗ | ✓ |
| Separation of tonic and phasic activity of the skin conductance response | ✗ | ✗ | ✓ |
| Inclusion of trials reinforced by electrical stimulation in absence of MR recordings | ✗ | ✗ | ✓ |
| Removal of trials in which no knowledge of the current contingencies is possible | ✗ | ✗ | ✓ |
| ANOVA: 2x2x2 (stage x stimulus x phase) or 2x2x2x2 after adding a group factor / 2x2x2x2x2 after adding group and contingency reversal factors when comparing patients with OCD and healthy controls | ✓ | ✗ | ✓ |
| Post-experiment questions verifying whether participants were aware of contingency reversal | ✗ | ✓ | ✓ |

**Supplementary Table 2.**
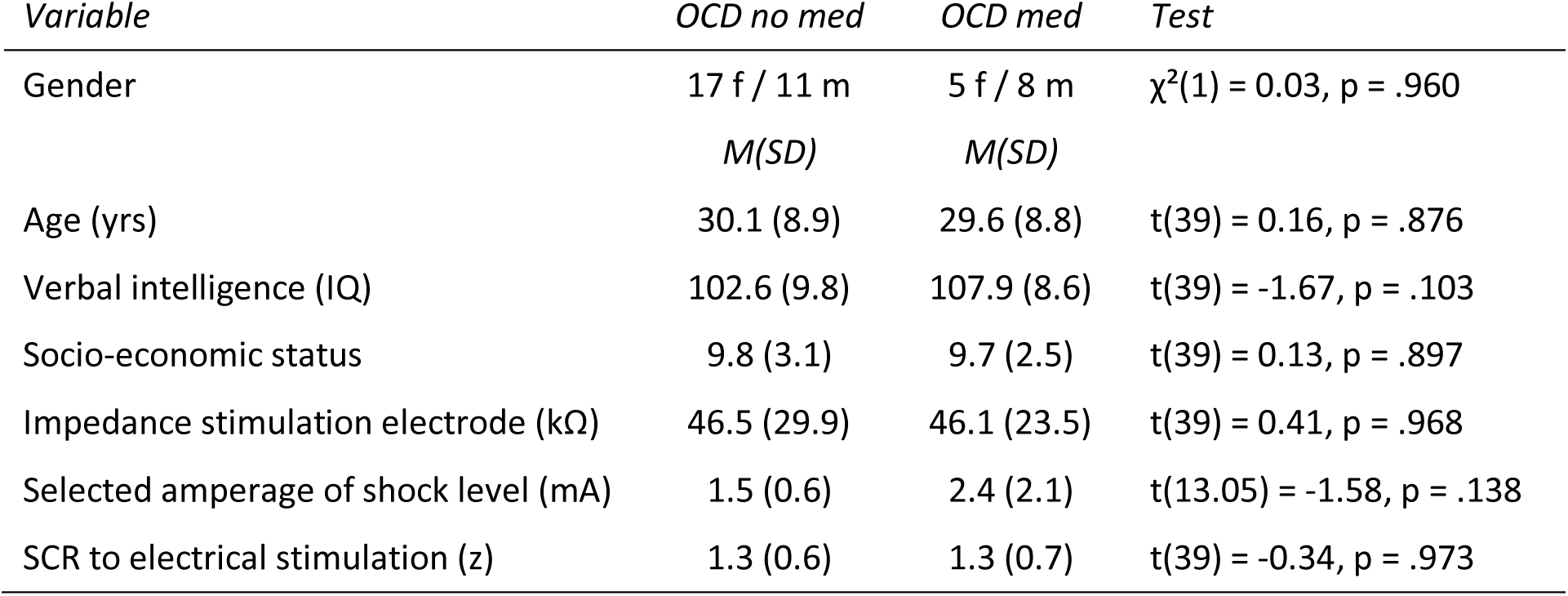
Group differences in demographic and basic experimental variables for the comparison of unmedicated and medicated patients.

**Supplementary Table 3.** ANOVA with repeated measures for unmedicated vs. medicated patients

| | F (1,37) | p | partial $\eta^2$ |
| --- | --- | --- | --- |
| ***Replication of fear reversal effects*** |  |  |  |
| Stage (acquisition vs. reversal) | = 5.47 | = .025 | .129 |
| Phase (early vs. late) | = 0.05 | = .825 | .001 |
| Stimulus (CS+ vs. CS-) | = 18.31 | < .001 | .331 |
| Stage x phase | = 0.11 | = .747 | .003 |
| Stage x stimulus | = 1.41 | = .243 | .037 |
| Phase x stimulus | = 2.27 | = .140 | .058 |
| Stage x phase x stimulus | = 0.01 | = .922 | .000 |
| ***Medication differences*** |  |  |  |
| Medication (unmedicated vs. medicated) | = 0.21 | = .653 | .006 |
| Medication x stage | = 0.02 | = .894 | .000 |
| Medication x phase | = 0.43 | = .514 | .012 |
| Medication x stimulus | = 0.01 | = .909 | .000 |
| Medication x stage x phase | = 2.11 | = .155 | .054 |
| Medication x stage x stimulus | = 1.19 | = .283 | .031 |
| Medication x phase x stimulus | = 0.17 | = .682 | .005 |
| Medication x stage x phase x stimulus | = 0.26 | = .613 | .007 |
| ***Reversal awareness differences*** |  |  |  |
| Awareness (aware vs. unaware) | = 2.04 | = .161 | .052 |
| Awareness x medication | = 2.48 | = .124 | .063 |
| Awareness x stage | = 0.68 | = .414 | .018 |
| Awareness x stage x medication | = 2.66 | = .111 | .067 |
| Awareness x phase | = 0.66 | = .422 | .017 |
| Awareness x phase x medication | = 0.53 | = .472 | .014 |
| Awareness x stimulus | = 1.68 | = .203 | .043 |
| Awareness x stimulus x medication | = 0.05 | = .818 | .001 |
| Awareness x stage x phase | = 1.28 | = .266 | .033 |
| Awareness x stage x phase x medication | = 0.67 | = .418 | .018 |
| Awareness x stage x stimulus | = 3.49 | = .070 | .086 |
| Awareness x stage x stimulus x medication | = 0.22 | = .639 | .006 |
| Awareness x phase x stimulus | = 0.00 | = .964 | .000 |
| Awareness x phase x stimulus x<br>medication | = 0.26 | = .613 | .007 |
| Awareness x stage x phase x stimulus | = 0.05 | = .819 | .001 |
| Awareness x stage x phase x stimulus x<br>medication | = 0.01 | = .925 | .000 |

## References

Apergis-Schoute, A. M., Gillan, C. M., Fineberg, N. A., Fernandez-Egea, E., Sahakian, B. J., & Robbins, T. W. (2017). Neural basis of impaired safety signaling in Obsessive Compulsive Disorder. Proc Natl Acad Sci U S A, 114(12), 3216–3221. doi: 10.1073/pnas.1609194114

Beck, A. T., Steer, R. A., & Brown, G. K. (1996). Beck depression inventory-II. San Antonio: The Psychological Corporation.

Benedek, M., & Kaernbach, C. (2010). A continuous measure of phasic electrodermal activity. Journal of neuroscience methods, 190(1), 80–91. doi: 10.1016/j.jneumeth.2010.04.028

Boedhoe, P. S., Schmaal, L., Abe, Y., Ameis, S. H., Arnold, P. D., Batistuzzo, M. C., … van den Heuvel, O. A. (2017). Distinct Subcortical Volume Alterations in Pediatric and Adult OCD: A Worldwide Meta- and Mega-Analysis. Am J Psychiatry, 174(1), 60–69. doi: 10.1176/appi.ajp.2016.16020201

Bui, E., Orr, S. P., Jacoby, R. J., Keshaviah, A., LeBlanc, N. J., Milad, M. R., … Simon, N. M. (2013). Two weeks of pretreatment with escitalopram facilitates extinction learning in healthy individuals. Hum Psychopharmacol, 28(5), 447–456. doi: 10.1002/hup.2330

Burghardt, N. S., Sigurdsson, T., Gorman, J. M., McEwen, B. S., & LeDoux, J. E. (2013). Chronic antidepressant treatment impairs the acquisition of fear extinction. Biol Psychiatry, 73(11), 1078–1086. doi: 10.1016/j.biopsych.2012.10.012

Burghardt, N. S., Sullivan, G. M., McEwen, B. S., Gorman, J. M., & LeDoux, J. E. (2004). The selective serotonin reuptake inhibitor citalopram increases fear after acute treatment but reduces fear with chronic treatment: a comparison with tianeptine. Biol Psychiatry, 55(12), 1171–1178. doi: 10.1016/j.biopsych.2004.02.029

Chamberlain, S. R., Menzies, L., Hampshire, A., Suckling, J., Fineberg, N. A., del Campo, N., … Sahakian, B. J. (2008). Orbitofrontal dysfunction in patients with obsessive-compulsive disorder and their unaffected relatives. Science, 321(5887), 421–422. doi: 10.1126/science.1154433

Cohen, R. A. (1993). The Orienting Response The Neuropsychology of Attention (pp. 95–113). Boston, MA: Springer.

Dieterich, R., Endrass, T., & Kathmann, N. (2017). Uncertainty increases neural indices of attention in obsessive-compulsive disorder. Depress Anxiety, 34(11), 1018–1028. doi: 10.1002/da.22655

Duits, P., Cath, D. C., Lissek, S., Hox, J. J., Hamm, A. O., Engelhard, I. M., … Baas, J. M. (2015). Updated meta-analysis of classical fear conditioning in the anxiety disorders. Depress Anxiety, 32(4), 239–253. doi: 10.1002/da.22353

Ekman, P., & Friesen, W. (1976). Pictures of facial affect. 1976. Palo Alto, CA: Consulting Psychologists.

Elsner, B., Wolfsberger, F., Srp, J., Windsheimer, A., Becker, L., Jacobi, T., … Reuter, B. (2020). Long-Term Stability of Benefits of Cognitive Behavioral Therapy for Obsessive Compulsive Disorder Depends on Symptom Remission During Treatment. Clinical Psychology in Europe, 2(1), 1–18. doi: 10.32872/cpe.v2i1.2785

Foa, E. B., Huppert, J. D., Leiberg, S., Langner, R., Kichic, R., Hajcak, G., & Salkovskis, P. M. (2002). The Obsessive-Compulsive Inventory: development and validation of a short version. Psychol Assess, 14(4), 485–496. doi: 10.1037/1040-3590.14.4.485

Geller, D. A., McGuire, J. F., Orr, S. P., Small, B. J., Murphy, T. K., Trainor, K., … Storch, E. A. (2019). Fear extinction learning as a predictor of response to cognitive behavioral therapy for pediatric obsessive compulsive disorder. J Anxiety Disord, 64, 1–8. doi: 10.1016/j.janxdis.2019.02.005

Goodman, W. K., Price, L. H., Rasmussen, S. A., Mazure, C., Delgado, P., Heninger, G. R., & Charney, D. S. (1989a). The Yale-Brown Obsessive Compulsive Scale. II. Validity. Archives of General Psychiatry, 46(11), 1012–1016. doi: 10.1001/archpsyc.1989.01810110054008

Goodman, W. K., Price, L. H., Rasmussen, S. A., Mazure, C., Fleischmann, R. L., Hill, C. L., … Charney, D. S. (1989b). The Yale-Brown Obsessive Compulsive Scale. I. Development, use, and reliability. Archives of General Psychiatry, 46(11), 1006–1011. doi: 10.1001/archpsyc.1989.01810110048007

Hand, I., & Büttner-Westphal, H. (1991). Die Yale-Brown Obsessive Compulsive scale (Y-BOCS): Ein halbstrukturiertes Interview zur Beurteilung des Schweregrades von Denk-und Handlungszwängen. Verhaltenstherapie, 1(3), 223–225. doi: 10.1159/000257972

Hansen, B., Kvale, G., Hagen, K., Havnen, A., & Öst, L. G. (2018). The Bergen 4-day treatment for OCD: four years follow-up of concentrated ERP in a clinical mental health setting. Cogn Behav Ther, 1–17. doi: 10.1080/16506073.2018.1478447

Hensman, R., Guimarães, F. S., Wang, M., & Deakin, J. F. (1991). Effects of ritanserin on aversive classical conditioning in humans. Psychopharmacology (Berl), 104(2), 220–224. doi: 10.1007/bf02244182

Hindi Attar, C., Finckh, B., & Büchel, C. (2012). The influence of serotonin on fear learning. PLoS One, 7(8), e42397. doi: 10.1371/journal.pone.0042397

Kaczkurkin, A. N., & Lissek, S. (2013). Generalization of Conditioned Fear and Obsessive-Compulsive Traits. J Psychol Psychother, 7, 3. doi: 10.4172/2161-0487.S7-003

Kanen, J. W., Apergis-Schoute, A. M., Yellowlees, R., Arntz, F. E., van der Flier, F. E., Price, A., … Robbins, T. W. (2020). Serotonin depletion impairs both Pavlovian and instrumental reversal learning in healthy humans. bioRxiv, 2020.2004.2026.062463. doi: 10.1101/2020.04.26.062463

Lampert, T., & Kroll, L. E. (2009). Die Messung des sozioökonomischen Status in sozialepidemiologischen Studien Gesundheitliche Ungleichheit (pp. 309–334): Springer.

Lissek, S., Kaczkurkin, A. N., Rabin, S., Geraci, M., Pine, D. S., & Grillon, C. (2014). Generalized anxiety disorder is associated with overgeneralization of classically conditioned fear. Biol Psychiatry, 75(11), 909–915. doi: 10.1016/j.biopsych.2013.07.025

Lissek, S., Rabin, S., Heller, R. E., Lukenbaugh, D., Geraci, M., Pine, D. S., & Grillon, C. (2010). Overgeneralization of conditioned fear as a pathogenic marker of panic disorder. Am J Psychiatry, 167(1), 47–55. doi: 10.1176/appi.ajp.2009.09030410

Lonsdorf, T. B., Klingelhöfer-Jens, M., Andreatta, M., Beckers, T., Chalkia, A., Gerlicher, A., … Merz, C. J. (2019). Navigating the garden of forking paths for data exclusions in fear conditioning research. Elife, 8. doi: 10.7554/eLife.52465

Mataix-Cols, D., Fernandez de la Cruz, L., & Rück, C. (2019). When Improving Symptoms Is Not Enough-Is It Time for Next-Generation Interventions for Obsessive-Compulsive Disorder? JAMA Psychiatry. doi: 10.1001/jamapsychiatry.2019.2335

Mataix-Cols, D., Rosario-Campos, M. C., & Leckman, J. F. (2005). A multidimensional model of obsessive-compulsive disorder. Am J Psychiatry, 162(2), 228–238. doi: 10.1176/appi.ajp.162.2.228

McLaughlin, N. C., Strong, D., Abrantes, A., Garnaat, S., Cerny, A., O’Connell, C., … Greenberg, B. D. (2015). Extinction retention and fear renewal in a lifetime obsessive-compulsive disorder sample. Behav Brain Res, 280, 72–77. doi: 10.1016/j.bbr.2014.11.011

Milad, M. R., Furtak, S. C., Greenberg, J. L., Keshaviah, A., Im, J. J., Falkenstein, M. J., … Wilhelm, S. (2013). Deficits in conditioned fear extinction in obsessive-compulsive disorder and neurobiological changes in the fear circuit. JAMA Psychiatry, 70(6), 608-618; quiz 554. doi: 10.1001/jamapsychiatry.2013.914

Mineka, S., & Oehlberg, K. (2008). The relevance of recent developments in classical conditioning to understanding the etiology and maintenance of anxiety disorders. Acta Psychol (Amst), 127(3), 567–580. doi: 10.1016/j.actpsy.2007.11.007

Montgomery, S. A., & Åsberg, M. (1979). A new depression scale designed to be sensitive to change. The British journal of psychiatry, 134(4), 382–389. doi: 10.1192/bjp.134.4.382

Mowrer, O. H. (1939). A stimulus-response analysis of anxiety and its role as a reinforcing agent. Psychological review, 46(6), 553–565. doi: 10.1037/h0054288

Nanbu, M., Kurayama, T., Nakazawa, K., Matsuzawa, D., Komiya, Z., Haraguchi, T., … Shimizu, E. (2010). Impaired P50 suppression in fear extinction in obsessive-compulsive disorder. Prog Neuropsychopharmacol Biol Psychiatry, 34(2), 317–322. doi: 10.1016/j.pnpbp.2009.12.005

Olatunji, B. O., Davis, M. L., Powers, M. B., & Smits, J. A. (2013). Cognitive-behavioral therapy for obsessive-compulsive disorder: a meta-analysis of treatment outcome and moderators. J Psychiatr Res, 47(1), 33–41. doi: 10.1016/j.jpsychires.2012.08.020

Öst, L. G., Havnen, A., Hansen, B., & Kvale, G. (2015). Cognitive behavioral treatments of obsessive-compulsive disorder. A systematic review and meta-analysis of studies published 1993-2014. Clin Psychol Rev, 40, 156–169. doi: 10.1016/j.cpr.2015.06.003

Remijnse, P. L., Nielen, M. M., van Balkom, A. J., Cath, D. C., van Oppen, P., Uylings, H. B., & Veltman, D. J. (2006). Reduced orbitofrontal-striatal activity on a reversal learning task in obsessive-compulsive disorder. Arch Gen Psychiatry, 63(11), 1225–1236. doi: 10.1001/archpsyc.63.11.1225

Schiller, D., Levy, I., Niv, Y., LeDoux, J. E., & Phelps, E. A. (2008). From fear to safety and back: reversal of fear in the human brain. J Neurosci, 28(45), 11517–11525. doi: 10.1523/jneurosci.2265-08.2008

Schmidt, K., & Metzler, P. (1992). Wortschatztest (WST). Beltz: Weinheim.

Spielberger, C., Gorsuch, R. L., & Lushene, R. (1970). STAI manual for the State-Trait Inventory. CA: Palo Alto.

Taylor, S., Abramowitz, J. S., & McKay, D. (2012). Non-adherence and non-response in the treatment of anxiety disorders. J Anxiety Disord, 26(5), 583–589. doi: 10.1016/j.janxdis.2012.02.010

Valerius, G., Lumpp, A., Kuelz, A. K., Freyer, T., & Voderholzer, U. (2008). Reversal learning as a neuropsychological indicator for the neuropathology of obsessive compulsive disorder? A behavioral study. J Neuropsychiatry Clin Neurosci, 20(2), 210–218. doi: 10.1176/appi.neuropsych.20.2.210

